# Integrative taxonomy and distribution of *Apis* species in Malaysia

**DOI:** 10.64898/2025.12.18.695282

**Authors:** Asha Devi Pallujam, Wai Leng Lee, Philipp Engel, Sze Huei Yek

## Abstract

Southeast Asia is the centre of diversification for honey bees (genus *Apis*). Hence, accurate identification of honey bees found here is instrumental in paving the way for studying evolutionary relationships, biogeographic patterns, and management of these key pollinators. In this study, we characterized *Apis* species diversity across peninsular Malaysia and Borneo Malaysia using an integrative approach combining morphological traits and mitochondrial markers (CO1 and 16S rRNA). Field collections included both managed and wild colonies of *A. mellifera, A. cerana, A. dorsata, A. florea,* and *A. andreniformis* from peninsular Malaysia, as well as two additional cavity-nesting species of Borneo *A. koschevnikovi* and *A. nuluensis*. Species identification was performed based on morphological features, nesting ecology, and subsequently validated through sequencing. Samples collected from peninsular Malaysia showed congruence between field/morphological and molecular identification. In contrast, cavity-nesting bees from Borneo exhibited overlapping morphological characteristics leading to misidentification of species. These findings suggest the reliability of carrying out field-based identification of honey bees in peninsular Malaysia but requires molecular validation for Borneo region. The limitations of morphology-based interpretation for Borneo cavity-nesting bees highlights the need for more population-level and speciation studies on these species.

## Introduction

Honey bees (genus *Apis*) are important social insects that provide pollination services for a wide variety of plants and produce valuable apiarian products such as honey, beeswax, propolis, and royal jelly (Crane 2009; Partap 2011; Iryani and Ismail 2016; Hung et al. 2018). The genus includes ten species, which are categorized into three distinct clusters: cavity-nesting bees (*A. mellifera*, *A. cerana*, *A. koschevnikovi*, *A. nuluensis*, and *A. nigrocincta*), giant honey bees (*A. dorsata*, *A. binghami* and *A. laboriosa*), and dwarf honey bees (*A. florea* and *A. andreniformis*). These species differ predominantly in nesting ecology, colony and body sizes, behaviour, and ecological adaptations (Willis et al. 1992; Tanaka et al. 2001; Arias and Sheppard 2005; Raffiudin and Crozier 2007). However, they all play vital roles through pollination of wild and cultivated plants (Crane 2009; Partap 2011). Among these species, *A. mellifera* has achieved global success due to its adaptability for honey production and pollination (Moritz et al. 2005), while other species are confined to Asia, often overlapping in tropical regions (Engel 1999; Arias and Sheppard 2005).

Among the five cavity-nesting bees, four species can be found in Malaysia - *A. mellifera*, *A. cerana*, *A. koschevnikovi* and *A. nuluensis* (Phiancharoen et al. 2011; Hepburn and Adloff 2011; Yang et al. 2011; Silva et al. 2020). They typically form their nests in enclosed spaces such as tree cavities or man-made hives (Phiancharoen et al. 2011; Gupta 2014). The nests are built of multiple combs, parallel to each other, with a uniform spacing in between the combs called ‘bee space’. The Western honey bee, *A. mellifera*, native to Africa, Europe and the Middle East, has been introduced to Asian countries, including Malaysia since the 1900s for honey production in the apiculture industry (Moritz et al. 2005; Iryani and Ismail 2016). In Malaysia, *A. mellifera* is primarily managed in apiaries and is rarely found in natural environments. This limits its direct competition for nesting cavities with native species, although it still competes for floral resources in the region (Iryani and Ismail 2016). The Asian honey bee, *A. cerana* is native to Southern and Southeastern Asia and plays a vital role in natural ecosystems and traditional beekeeping practices. It is well adapted to tropical climates and is resilient to local pests and diseases, making it an important species for small-scale apiculture (Phiancharoen et al. 2011). The red honey bee, *A. koschevnikovi,* is endemic to Southeast Asia and has a restricted distribution, primarily found in lowland rainforests. It is sympatric with two other cavity-nesting bees (*A. mellifera* and *A. cerana*) but is considerably less common. Despite its restricted distribution, its ecological role in pollinating forest plants is crucial for maintaining biodiversity in these habitats (Hepburn and Adloff 2011). Unlike *A. mellifera* and *A. cerana, A. koschevnikovi* has not been widely utilized in apiculture, largely due to its limited range and forest habitat requirements. The Borneo highland honey bee, *A. nuluensis*, is endemic to 1900 m asl at Kinabalu and Crocker Range of Sabah, Borneo Malaysia (Tingek et al. 1996; Yek et al. 2024). This species is closely related to *A. cerana* but has adapted to highland living (Tanaka et al. 2001). Because of its limited and restricted distribution in the protected highland area, *A. nuluensis* has not been domesticated in apiculture.

The giant honey bee, *A. dorsata* is a migratory species with a wide distribution across Southeast Asia, including peninsular Malaysia. *A. dorsata* builds exposed, large single-comb (up to 1 m) nests on tall trees or cliffs, usually from 3 to 25 m above the ground (Gupta 2014). Nests of *A. dorsata* may occur singly or in groups in a single tall tree called a ‘bee tree’. *A. dorsata* is significant for its role in forest pollination and as a source of wild honey. This species exhibits remarkable foraging abilities, often covering long distances to collect nectar and pollen; and they have the ability to fly at night when the moonlight is adequate (Crane 1999; Gupta 2014; Iryani and Ismail, 2016). *A. dorsata* exhibits an annual migratory behaviour, travelling 100-200 km during seasonal transitions from rainy to dry periods to adapt to fluctuations in resource availability and environmental conditions (Robinson 2012).

The two dwarf honey bees, *A. florea* and *A. andreniformis,* are sister species with a partially sympatric distribution in southern Asia. They build small, exposed nests on shrubs or on low hanging tree branches, and their foraging range is quite limited - often remaining within 750 m of the nest (Crane 2009; Hepburn and Adloff 2011; Gupta 2014).

*A. florea* is native to southern Asia and parts of the Middle East; and has been expanding its range into East Africa, Southeast Asia and East Asia (Silva et al. 2020; Ascher et al. 2022). *A. florea,* a non-native species in Malaysia, has successfully established colonies in peninsular Malaysia but not in Borneo, Malaysia (Silva et al. 2020). However, information on its establishment history and ecological impacts remains limited, indicating a need to investigate further. *A. andreniformis* can be found naturally from the foothills of the eastern Himalayas towards Indochina, Sundaland and the Philippines, including Malaysia. While *A. florea* is adapted to more seasonal and open vegetation areas, *A. andreniformis* tends to dominate in mesic and densely forested habitats (Oldroyd and Wongsiri 2006; Hepburn and Adloff 2011; Silva et al. 2020).

The phylogenetic relationship among *Apis* species have been widely studied using nuclear and mitochondrial DNA markers such as ND2, EF1-ALPHA, 16S rRNA, COI, and COII (Willis et al. 1992; Tanaka et al. 2001; Arias and Sheppard 2005; Raffiudin and Crozier 2007; Kek et al. 2017; Khan 2021). While molecular tools such as DNA barcoding have been increasingly applied for species identification, their consistent use across *Apis* species in this region is lacking. Recent reports indicating the expansion of *A. florea* into the region highlight the need for comprehensive molecular and ecological studies to confirm its presence and understand its interactions with other native species (Silva et al. 2020; Ascher et al. 2022).

Although morphological identification methods remain valuable, they often fall short in discriminating between closely related species, particularly among cavity-nesting species due to their overlapping morphological and behavioural characteristics traits. To address these challenges, this study characterizes *Apis* species diversity across peninsular Malaysia and Borneo Malaysia using an integrative approach that combines morpho-taxonomic assessment with mitochondrial markers (CO1 and 16S rRNA). The identification of species was first performed based on morphological features and nesting ecology and subsequently validated through molecular sequencing to ensure accurate delineation of taxa. Our results show that morphological identification is largely reliable for the *Apis* species of peninsular Malaysia, whereas closely related cavity-nesting species in Borneo require molecular confirmation due to overlapping traits and frequent misidentification.

## Materials and method

### Ethics statement

This research involved collecting honey bees from forest reserves and national parks. For peninsular Malaysia samples, the permit application was approved by the Perak State Forestry Department to collect the samples from Temenggor forest reserve with the approval number: JH/100 JLD.31 (46). Access and benefit sharing (ABS) was also applied with the application reference number: 676216. For Borneo samples, the permit application was approved by the Sabah Biodiversity Centre, with the access license JKM/MBS.1000-212 JLD.16 (117), and Sabah Parks approval TTS 100-6/2 JLD.34 issued to SHY (Supplementary Information – Permits, Sample Collection).

### Sample collection

Extensive sampling of honey bees was conducted across peninsular Malaysia from December 2021 to October 2022. Borneo honey bee sampling was carried out from November 2023 to February 2024. Worker bees were either collected from the nest entrance or from flowers (Table 1; Supplementary Table S1). Out of the four cavity-nesting bees, the Western honey bee, *A. mellifera,* was collected only from managed colonies, while the native species, *A. cerana* was collected from both managed as well as wild colonies. No colonies of *A. koschevnikovi* were detected during the field surveys from the peninsular Malaysia sampling, despite recorded occurrence of this species in the region. *A. koschevnvikovi* and *A. nuluensis* were only collected from wild colonies, in national parks and university forest fragments in Borneo Malaysia. The giant honey bee *A. dorsata,* and dwarf honey bees (*A. florea* and *A. andreniformis*) were also collected from wild colonies. Most of the wild species’ collection was carried out with the assistance of a local bee relocation Non-Governmental Organisation called MY Bee Saviour in peninsular Malaysia, whereas the wild species’ collection in Borneo was carried out with the assistance of Sabah Parks nature guides.

**Table 1:** List of collected samples, the corresponding sampling sites and the GenBank accession numbers of the sequences of two analyzed genes (CO1 and 16S rRNA).

| Sample ID | Location | Coordinates | Field<br>identification | Molecular<br>identification | GenBank accession numbers |  |
| --- | --- | --- | --- | --- | --- | --- |
|  |  |  |  |  | CO1 | 16S rRNA |
| A.mellifera_1 | Sitiawan, Perak | N04°14.435'<br>E100°39.484' | *A. mellifera* | *A. mellifera* | OR563685 | OR514819 |
| A.mellifera_2 | Sitiawan, Perak | N03°14.622'<br>E100°39.516' | *A. mellifera* | *A. mellifera* | OR563672 | OR514815 |
| A.mellifera_3 | Sitiawan, Perak | N04°16.380'<br>E100°37.378' | *A. mellifera* | *A. mellifera* | OR563671 | OR514814 |
| A.mellifera_4 | Baling, Kedah | N05°35.670'<br>E100°46.794' | *A. mellifera* | *A. mellifera* | OR563670 | OR514807 |
| A.mellifera_5 | Cameron Highlands,<br>Perak | N04°26.826'<br>E101°23.565' | *A. mellifera* | *A. mellifera* | OR563667 | OR514803 |
| A.mellifera_6 | Cyberjaya, Selangor | N02°57.019'<br>E101°37.936' | *A. mellifera* | *A. mellifera* | OR563660 | OR514795 |
| A.cerana_1 | Monash University,<br>Selangor | N 03°03.843'<br>E101°36.088' | *A. cerana* | *A. cerana* | OR563681 | OR514820 |
| A.cerana_2 | Cheras, Selangor | N03°02.924'<br>E101°44.310' | *A. cerana* | *A. cerana* | OR563678 | OR514817 |
| A.cerana_3 | Kepala Batas, Penang | N05°30.539'<br>E100°25.858' | *A. cerana* | *A. cerana* | OR563669 | OR514813 |
| A.cerana_4 | Kepala Batas, Penang | N05°30.541'<br>E100°25.860' | *A. cerana* | *A. cerana* | OR563668 | OR514805 |
| A.cerana_5 | Cameron Highlands,<br>Perak | N04°26.826'<br>E101°23.566' | *A. cerana* | *A. cerana* | OR563666 | OR514804 |
| A.cerana_6 | MPSJ, Selangor | N03°02.848'<br>E101°34.188' | *A. cerana* | *A. cerana* | OR563664 | OR514801 |
| A.dorsata_1 | Temenggor Forest,<br>Perak | N05°27.402'<br>E101°21.611' | *A. dorsata* | *A. dorsata* | OR563675 | OR514816 |
| A.dorsata_2 | Temenggor Forest,<br>Perak | N05°25.014'<br>E101°21.828' | *A. dorsata* | *A. dorsata* | OR563675 | OR514810 |
| A.dorsata_3 | Temenggor Forest,<br>Perak | N05°28.514'<br>E101°21.754' | *A. dorsata* | *A. dorsata* | OR563674 | OR514809 |
| A.dorsata_4 | Gerik, Perak | N05°29.758'<br>E101°11.285' | *A. dorsata* | *A. dorsata* | OR563673 | OR514808 |
| A.dorsata_5 | Petaling Jaya,<br>Selangor | N04°26.826'<br>E101°23.566' | *A. dorsata* | *A. dorsata* | OR563665 | OR514802 |
| A.dorsata_6 | Shah Alam, Selangor | N03°06.151'<br>E101°28.582' | *A. dorsata* | *A. dorsata* | OR563663 | OR514800 |
| A.florea_1 | Petaling Jaya,<br>Selangor | N03°03.227'<br>E101°34.660' | *A. florea* | *A. florea* | OR563680 | OR514812 |
| A.florea_2 | Balik Pulau, Penang | N05°22.218'<br>E100°13.243' | *A. florea* | *A. florea* | OR563679 | OR514806 |
| A.florea_3 | Kajang, Selangor | N02°59.190'<br>E101°45.317' | *A. florea* | *A. florea* | OR563661 | OR514797 |
| A.florea_4 | Johor Bahru, Johor | N01°28.784'<br>E103°41.189' | *A. florea* | *A. florea* | OR563686 | OR514821 |
| A.florea_5 | Taman Kota<br>Laksamana, Melacca | N02°11.735'<br>E102°13.882' | *A. florea* | *A. florea* | OR563687 | OR514822 |
| A.florea_6 | Seremban, Negeri<br>Sembilan | N01°41.798'<br>E101°53.934' | *A. florea* | *A. florea* | OR563688 | OR514823 |
| A.andreniformis_1 | Cheras, Selangor | N03°02.974'<br>E101°44.333' | *A. andreniformis* | *A. andreniformis* | OR563682 | OR514818 |
| A.andreniformis_2 | Cheras, Selangor | N03°02.837'<br>E101°39.429' | *A. andreniformis* | *A. andreniformis* | OR563683 | OR563696 |
| A.andreniformis_3 | Mutiara Seputeh,<br>Kuala Lumpur | N03°06.938'<br>E101°41.342' | *A. andreniformis* | *A. andreniformis* | OR563677 | OR514811 |
| A.andreniformis_4 | Shah Alam, Selangor | N03°06.151'<br>E101°28.582' | *A. andreniformis* | *A. andreniformis* | OR563684 | OR514799 |
| A.andreniformis_5 | Shah Alam, Selangor | N03°06.821'<br>E101°28.559' | *A. andreniformis* | *A. andreniformis* | OR563662 | OR514798 |
| A.andreniformis_6 | Negeri Sembilan,<br>Seremban | N02°41.179'<br>E101°53.884' | *A. andreniformis* | *A. andreniformis* | N/A | OR514796 |
| A.cerana_7 | Poring Village,<br>Sabah | N06°02.503'<br>E116°42.374' | *A. cerana* | *A. cerana* | PX463715 | PX242781 |
| A.cerana_8 | UMS, Sabah | N06°02.3554'<br>E116°06.9248' | *A. cerana* | *A. cerana* | PX463700 | PX242782 |
| A.cerana_9* | Tahubung, Sabah | N06°03.00.2617'<br>E116°32.4962' | *A. nuluensis* | *A. cerana* | PX463722 | PX242783 |
| A.cerana_10* | Kiau Village, Sabah | N06°03.03.2084'<br>E116°30.1676' | *A. nuluensis* | *A. cerana* | PX463705 | PX242784 |
| A.cerana_11 | Kiau Village, Sabah | N06°03.03.2300'<br>E116°30.1975' | *A. cerana* | *A. cerana* | PX463723 | PX242785 |
| A.koschevnikovi_1 | UMS, Sabah | N06°02.3554'<br>E116°06.9248' | *A. koschevnikovi* | *A. koschevnikovi* | PX463718 | PX242786 |
| A.koschevnikovi_2* | Poring Village,<br>Sabah | N06°02.503'<br>E116°42.374' | *A. cerana* | *A. koschevnikovi* | PX463719 | PX242787 |
| A.nuluensis_1 | Mesilau, Sabah | N06°02.44.9'<br>E116°35.352' | *A. nuluensis* | *A. nuluensis* | PX463701 | PX242788 |
| A.nuluensis_2 | Mount Kinabalu,<br>Sabah | N06°03.2618'<br>E116°33.9178' | *A. nuluensis* | *A. nuluensis* | PX463721 | PX242789 |

\*Denotes samples with inconsistent field/morphology-based and molecular-based
identifications. All inconsistent samples were from Borneo Malaysia.

Four colonies of *A. dorsata* were collected from the Belum-Temenggor Tropical Rainforest in March 2022 where the bees built single nest comb under the major branches of emergent dipterocarp trees *Koompassia excelsa* (local name: Tualang) and fig tree, *Ficus alba* (local name: Pokok Ara). Since the bees build their comb about 30 m above the ground level, the honey harvesters that usually climb up to harvest honey were hired to assist in sample collection. Two colonies of *A. dorsata* were collected from urban areas. The dwarf honey bee species, *A. florea* and *A. andreniformis,* were collected from semi-urban areas in thick shrubs. The collection of dwarf honey bees was performed based on accessibility, with sites located by MY Bee Saviour. In peninsular Malaysia, the MY Bee Saviour are called upon to relocate nests of *Apis* species found in urban areas. This means they mostly encounter *A. dorsata*, *A. florea* and *A. andreniformis*. These three *Apis* species can be easily distinguished by body size and nesting ecology. However, MY Bee Saviour volunteers are not active in Borneo, Malaysia. Hence, *Apis* species were only distinguished by where they nest (inside cavities, or hanging on tree branches), body sizes and colouration. The least sampled *A. koschevnikovi* and *A. nuluensis* nests can only be found in naturally formed tree cavities. All *A. koschevnikovi* and *A. nuluensis* nests were found either through extensive searching by author (SHY) or Sabah Parks nature guides.

Worker bees were sampled using either plastic jars or laundry bags. Samples were anaesthetised in 95% ethanol and transported to the laboratory for storage at -20°C before further processing. The sampling and handling protocol strictly followed the Standard Operating Protocol (SOP) developed in collaboration with Engel’s Lab, University of Lausanne, Switzerland (Supplementary Information – SOP, Sample collection). The preliminary field identification was carried out by MY Bee Saviour volunteers, and Sabah Parks nature guides. Back at the laboratory, the identification was again checked using the morphological traits identification guide <u>Apoidea species -- identification guide -- Discover Life</u>.

### Sample processing and DNA extraction

One bee was used for each colony to ensure a broad representation of the population. Under a stereomicroscope and using sterile dissection tools, the head and abdomen were removed to ensure that the sample was free from tissues with high microbial loads to avoid contamination. The thorax, which is rich in muscle tissue and a reliable source of high-quality genomic DNA, was then isolated. To facilitate efficient grinding and subsequent lysis, the thoracic cuticle was carefully removed, leaving only the underlying muscle tissue. The cleaned tissue was then cut into small pieces and transferred into a sterile microcentrifuge tube and processed for total genomic DNA extraction using DNeasy Blood and Tissue Kit (Qiagen) following the manufacturer’s protocol with minor modifications. Instead of the recommended lysis incubation time of 1-3 h, it was increased to four hours. The extracted DNA was quantified by BioDrop ulite (Biochrom) and qualitatively evaluated by gel electrophoresis.

### PCR amplification and sequencing

Two mitochondrial markers - CO1 and 16S rRNA genes were chosen for their proven efficacy in resolving species-level discriminations and confirming identification accuracy in honey bees (Ozdil 2009; Kek et al. 2017; Khan 2021). The CO1 gene is the standard DNA barcode to identify and differentiate insect species as it has discriminatory power for most insect groups (Ozdil 2009). The 16S rRNA gene, on the other hand, is among the most commonly used mitochondrial DNA markers in *Apis* phylogenetic analysis (Kek et al. 2017; Khan 2021). Together, these markers offer a robust framework for accurately identifying *Apis* species in Malaysia.

The extracted genomic DNA was PCR amplified for CO1 and 16S rRNA genes using gene-specific primers: LepF1 5’-ATTCAACCAATCATAAAGATATTGG-3’; and LepR1 5’-TAAACTTCTGGATGTCCAAAAAATCA-3’ (Hebert et al. 2004); and WLO001F-Af 5’-AGGTCGATCTGCTCCATGAA-3’; and WLO001R-Af 5’-TCAACATCGAGGTCGCAATCA-3’. The latter primers were designed for this study based on *A. florea* mitochondrial 16S rRNA partial gene segment (Accession: L22894.1) using Primer3 (https://www.primer3plus.com/).

The amplification of genes was carried out in a 25 μl total volume, containing 12.5 μl of One *Taq* 2X master mix (New Zealand Biolabs. Inc), 0.5 μl of each primer (10 uM), 1 μl of template DNA, and 10.5 μl of nuclease-free water. The PCR amplification for the CO1 gene was carried out at 94 °C/30 s for denaturation, 48 °C/60 s for annealing, and 68 °C/60 s for extension. The cycle was repeated 30 times followed by an additional extension step of 68 °C for five minutes. The PCR amplification for 16S rRNA gene was carried out similarly except for the annealing step at 52 °C for 60 s. The PCR products were separated by gel electrophoresis on a 1.2 % agarose TAE buffer, and the amplified bands were visualized using the Gel Doc EZ imaging system (BioRad). The PCR products were subsequently purified using PCR purification Kit (Qiagen) following the manufacturer’s protocol. The purified products were then outsourced to the Genomics facility for Sanger sequencing using Applied Biosystems 3730xl DNA Analyzer (Macrogen Asia Pacific, Singapore).

#### Sequence analysis and species identification

The nucleotide sequences obtained from Sanger sequencing for both gene targets were first subjected to quality control. Low-quality regions were manually trimmed using MEGA 11 (Tamura et al. 2021). One CO1 gene sequence (A. andreniformis_6) was excluded from further analysis due to poor quality, leaving the remaining 38 high-quality sequences for species identification. All the 16S rRNA gene dataset yielded high-quality sequences. The forward and reverse sequences were then aligned, and consensus sequences were generated with EMBOSS Cons (Rice et al. 2000). The species identities were examined using BLAST searches on the NCBI database. All sequence data from this study were submitted to GenBank and assigned their corresponding accession numbers (Table 1).

## Results

### Apis species field collection and identification

For peninsular Malaysia, six colonies per species were sampled from five honey bee species - *A. mellifera, A. cerana, A. dorsata, A. florea,* and *A. andreniformis* (Fig. 1). The preliminary identification of these bee species was carried out in the field by either the beekeeper, MY Bee Saviour volunteers, or the author (ADP). The bees’ body size and colour, nesting substrates (cavity-nesters or open-nesters) and shape of combs were the main characteristics used for preliminary field identification. The Western honey bee *A. mellifera* was found only in managed farms in rows of neatly packed combs (Fig. 1a-b). The Asian honey bee *A. cerana*, on the other hand, was found in farms, wall cavities and natural tree cavities (Fig. 1c). Both *A. mellifera* and *A. cerana* are cavity nesters, with *A. mellifera* workers being substantially larger (10-15 mm) compared to *A. cerana* (10-11 mm). *A. mellifera* was easily identified with the help of bee farmers. The native species, *A. cerana*, was collected from both managed as well as wild colonies. To differentiate *A. mellifera* and *A. cerana* in the laboratory, the bees’ hind wing venation was examined under a stereo microscope. The hind wing of *Apis cerana* has a distal abscissa, but this structure is absent in *A. mellifera* (<u>Apoidea species -- identification guide -- Discover Life</u>).

**Fig. 1.**
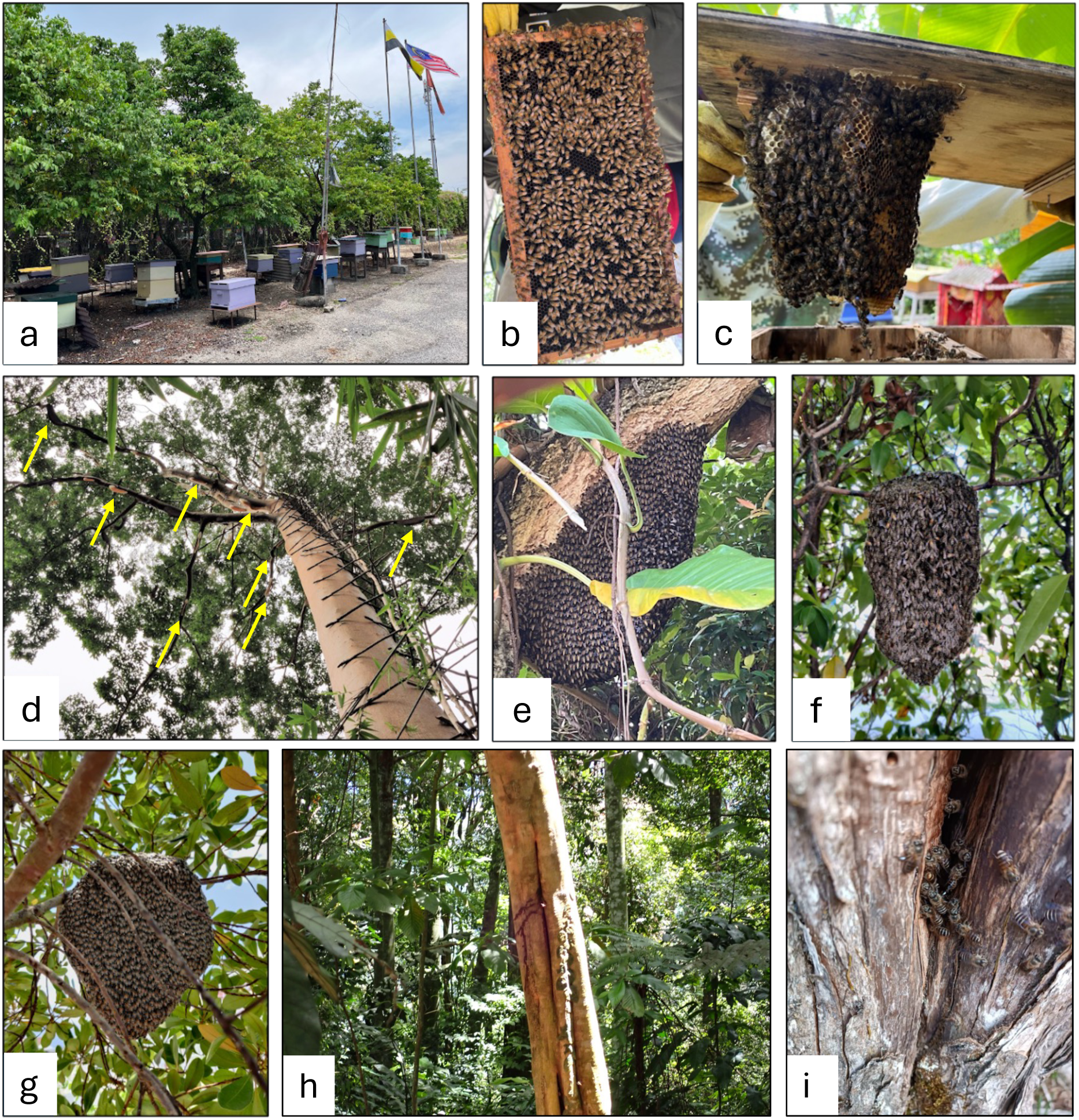
Nesting types of *Apis* species (a) Managed *A. mellifera* bee farm (b) Hive frame with bees from a managed *A. mellifera* colony (c) *A. cerana* colony inside a managed bee hive (d) Multiple nests of *A. dorsata* (yellow arrows) under the branches of *Koompassia excelsa* (Tualang tree) (e) A close-up image of an *A. dorsata* colony (f) A single comb of an *A. andreniformis* colony on a small branch of *Syzygium smithii* (Lilly Pilly) (g) *A. florea* nest on a small branch of an *Acacia* tree (h) *A. koschevnikovi* nest at Sabah Parks Kimanis substation (i) A cavity nest entrance of *A. nuluensis* at 3190 m above sea level at Mt. Kinabalu

The giant honey bee, *A. dorsata,* is mostly found in the lowland rainforest of peninsular Malaysia (Iryani and Ismail 2016). *A. dorsata* is an open nester and tends to build single-comb nests high up in the tree. For example, colonies collected from pristine forests were nesting approximately 30 m above ground on *Ficus alba* (Ara tree) and *Koompassia excelsa* (Tualang tree) (Fig. 1d), each comb measuring approximately 1.5 m x 1 m. Two out of the six colonies were collected from urban environments, where the nests were comparatively smaller and built approximately 15 m above ground (Fig. 1e). The colony sampled at Bongor village in Gerik, Perak State, was found nesting within creeper bushes with a comb measuring about 0.7 m x 0.6 m. At Setia Eco Park, Shah Alam, a single-comb nest was built on a *Tabebuia aurea* (Trumpet tree) branch approximately 15 m above ground with a comb measuring about 0.5 m x 0.4 m.

The dwarf honey bee species, *A. florea* and *A. andreniformis*, similar to the giant honey bee, are also open-nesters with single-comb nests. These bees were predominantly found nesting in semi-urban sites, such as community and household gardens. Both species appeared similar in overall body size and nest structure (Fig. 1f-g). Therefore, their colonies can only be distinguished upon closer inspection of the bees’ morphology under the stereo microscope. However, experienced field collectors can differentiate the species using differences in colouration, abdominal patterning, and body proportion (<u>Apoidea species -- identification guide -- Discover Life</u>). *A. florea* workers typically exhibited a lighter reddish-brown colouration while *A. andreniformis* is generally darker, with the worker’s first abdomen segment completely dark (Oldroyd and Wongsiri 2006). In some sites, colonies of *A. florea* and *A. andreniformis* were observed nesting only a few metres apart, on small trees in house gardens.

For Borneo Malaysia, the honey bee collection focuses on cavity-nesting *Apis* species that were not collected (*A. koschevnikovi*) or do not occur (*A. nuluensis*) in peninsular Malaysia (Fig. 1h-i). Although *A. koschevnikovi* was reported to be present in peninsular Malaysia, our one-year intensive sampling did not come across this species. This could be because *A. koschevnikovi* is rare and/or more commonly found in pristine forests in peninsular Malaysia. In Borneo Malaysia, *A. koschevnikovi* were relatively easily located as sampling was concentrated in forested habitats. All nests of *A. koschevnikovi* were found nesting inside hollow tree cavities in pristine forests and in lowland forests (∼500 m asl.). Since *A. koschevnikovi* co-occurs with *A. cerana*, it was not easy to distinguish between these two species in the field collection, because *A. cerana* is known to display variation in body size and colouration (Zhang et al. 2025). Even in the laboratory, examination of hind leg colour distribution (Koeniger et al. 2010) led to inaccurate identification, which was later corrected using molecular markers (see below).

The highland honey bee, *A. nuluensis,* is an endemic cavity-nesting species found only in Borneo Malaysia. This species can be found at elevations above 1600 m asl on the Kinabalu and Crocker Range mountains (Tingek et al. 1996). Foragers of *A. nuluensis* at the overlapping zone (1,600 to 1,900 m asl) with *A. koschenikovi* and *A. cerana* can sometimes lead to inaccurate identification due to the reasons mentioned above. However, from 1,900 m asl onwards, only *A. nuluensis* thrives (Yek et al. 2024). The highest densities of *A. nuluensis* nests were found between 1,900 and 3,200 m asl, on the Kinabalu Park mountains. They can nest in live or dead tree cavities (Supplementary Table S1).

### Molecular identification of *Apis* species

To confirm the preliminary field identification, one worker bee from each colony was used for molecular species identification. Both the CO1 and 16S rRNA gene fragments were successfully amplified by PCR, yielding the expected amplicon sizes of 680 bp and 424 bp, respectively (Supplementary Fig S1). Sequences with identity values between 99% and 100% relative to the NCBI database entries were considered definitive for species assignment, thereby validating the accuracy of species identification (Kek et al. 2017). All the field-based identifications of honey bee species from peninsular Malaysia were consistent with molecular classifications based on mitochondrial markers. However, molecular identification results indicated discrepancies between field identifications and sequence-based species assignment for Borneo Malaysia samples. This is particularly true among cavity-nesting species, such as *A. cerana, A. nuluensis,* and *A. koschevnikovi*. For example, *A. cerana* was mistaken as *A. nuluensis* in the overlapping zone (1,600 m asl) and *A. koschevnikovi* as *A. cerana* in the co-occurred elevation (∼500 m asl) (Table 1). Due to the variations displayed by *A. cerana*, field identification for these three cavity-nesting species requires caution.

## Discussion

The successful amplification of CO1 and 16S rRNA genes and their congruence with NCBI reference sequences confirm the reliability of these mitochondrial markers to delineate the species boundaries of honey bees (Ozdil 2009; Kek et al. 2017; Khan 2021). The field-based identifications of peninsular Malaysia are consistent with those obtained through molecular identification. However, the inconsistencies between field and molecular markers of some Borneo Malaysia samples highlight the need for caution in relying solely on morphological identification, especially for cavity-nesting bees (*A. cerana*, *A. koschenikovi*, and *A. nuluensis*).

In peninsular Malaysia, *A. cerana* populations, unlike *A. mellifera*, are predominantly feral and frequently occupy cavities in both natural and artificial structures. They can be found nesting in natural tree cavities, building crevices, and abandoned bee boxes. They often exhibit a transient nesting strategy, occupying cavities for a limited period before absconding, a well-known behaviour of native *A. cerana* populations (Pokhrel et al. 2006; Anna 2013). This flexible nesting behaviour allows *A. cerana* to exploit a wide range of habitats and to rapidly recolonise new sites following changes in the environment and resources. In contrast, *A. mellifera* is fully domesticated in the region, where it is maintained in managed apiaries for honey production and pollination services (Iryani and Ismail 2016). We have not encountered *A. mellifera* outside apiaries. Hence, *A. mellifera* is unlikely to compete directly with *A. cerana* or *A. koschenikovi* on nesting resources. However, they may still compete with native bee species for nectar and pollen resources.

*A. dorsata* is morphologically the largest among the *Apis* species found in the region and is known for constructing large, single-comb nests on the branches of tall trees in its natural forest habitat (Gupta 2014). This species is highly migratory and exhibits flexible nesting behaviour outside its natural habitat, adapting to changes in season and resource availability. The collections of smaller, transient nests in urban and peri-urban areas are consistent with the non-flowering seasons in their natal forest habitats (Robinson 2012; Robinson 2021). *Apis dorsata* colonies typically migrate during monsoon (November - March) from the forest to drier lowland when flower resources decline in the rainforest (personal communication with bee hunters). The bees return between April and May, aligning with the inter-monsoon period when the mass blooming of dipterocarps flowers starts. Such nesting patterns and long-distance migratory movements have been widely reported across the species’ distribution range in South and Southeast Asia (Phiancharoen et al. 2011; Robinson 2012; Gupta 2014).

*A. andreniformis* - a native dwarf honey bee species prefers shaded and forested habitats, while the introduced dwarf honey bee *A. florea* is known to thrive in open vegetation areas (Oldroyd and Wongsiri 2006; Hepburn and Adloff 2011). The occurrence of both species in proximity within semi-urban areas suggests a potential overlap in habitat use, likely facilitated by the availability of suitable nesting substrates and floral resources. This spatial overlap also indicates ecological flexibility and potential competition between the two dwarf honey bees (Silva et al. 2020; Hsu et al. 2022). The capacity of *A. florea* in range expansion and local population increases has generated concern about potential ecological impacts when it appears outside historical ranges. The potential of *A. florea* to act as a vector of pathogens warrants consideration, particularly in regions where its populations are recently established or expanding (El-Niweiri et al. 2019; Silva et al. 2020; Hsu et al. 2022). Future work should focus on long-term ecological monitoring of *A. florea* in peninsular Malaysia, as this species was collected in proximity to native *A. andreniformis*. Studies such as comparative assessments of floral resource partitioning, colony density, and pathogen load are required to elucidate the extent of ecological overlap and potential interspecific interactions among these two dwarf honey bee species.

Borneo constitutes a regional hotspot for cavity-nesting *Apis* diversity and harbours closely related taxa with subtle morphological differences. Mitochondrial and phylogeographic studies show deep structure within the Asian cavity-nesting clade and reveal distinct lineages that complicates simple field assignments (Takahashi et al. 2017; Su et al. 2023). Morphometric analyses showed that Bornean cavity-nesting species (*A. cerana, A. koschevnikovi,* and *A. nuluensis*) overlap in many conventional characteristics (body size, colouration, tomentum pattern, and wing venation indices) that could frequently result in inaccurate identification (Rinderer et al. 1989; Fuchs et al., 1996). Molecular studies have provided both confirmation of cryptic diversity (e.g., mitogenomes for *A. nuluensis* and *A. cerana* lineages) and practical diagnostic tools for delimiting taxa that are morphologically similar (Takahashi et al. 2017; Okuyama et al. 2017). Therefore, in regions where multiple cavity-nesting species are sympatric, reliance on field identification or casual morphological characters risk systematic misinterpretation and may bias studies on species distribution, niche partitioning, and pathogen dynamics.

We were unable to locate *A. koschevnikovi* nests despite extensive field collections over one year across various sites in peninsular Malaysia. The lack of samples for this species suggests its absence or an extremely limited distribution in the regions surveyed. From the literature, *A. koschevnikovi* is typically confined to tropical evergreen forests and has a potentially restricted range (Hadisoesilo et al. 2008). This species has also been reported to thrive in humid primary forests and their vicinity but is seldom found in disturbed or fragmented landscapes (Otis 1996). The collection of colonies for peninsular Malaysia primarily relied on reports from MY Bee Saviour volunteers and beekeepers, which focus on disturbed and urban fragmented landscapes. Hence, the absence of *A. koschevnikovi* samples in this study does not necessarily confirm its overall absence from peninsular Malaysia; rather, it underscores the need for further targeted surveys in forested habitats.

This study assesses the reliability of using known honey bee nesting ecology and morphological traits to identify honey bee species in Malaysia. We confirm the field and morphological identification with two molecular markers commonly used for bee species identification. Our results demonstrate that for honey bees in peninsular Malaysia, morphological traits, combined with field expertise from local beekeepers and organisations such as MY Bee Saviour, provide a dependable basis for species identification when molecular resources are limited. However, caution should be practised for differentiating cavity-nesting bee species (*A. cerana*, *A. koschenikovi*, and *A. nuluensis*) in Borneo Malaysia. We argued that at least one molecular marker (e.g., COI) should be used to confirm the identity of cavity-nesting bees sampled from Borneo, Malaysia.

## Acknowledgments

We thank the following organizations, bee farms, and people for their support in honey bee sampling and access - MY Bee Saviour, Nature Inspired (John Chan, Steven Wong, and JC Tan), Mr. Honeybee farm, Sherwani Honeybee, Mr. Wee, Steve Honey Bee farm, Cameron Highlands Bee Farm, Mr. Law, the Orang Asli community, and the Sabah parks nature guides for generously helping with the samples for this research. Thanks to Aiswarya Prasad, University of Lausanne, Switzerland, for helping with the sample collection standard protocol. We also thank Dr. Tan Hock Siew for his advice on the molecular work and Dr. Cyren Wong Zhi Hoong for helping with the morpho taxonomic analysis at Monash University Malaysia.

## Author Contributions

Conceptualization – SHY, ADP; Methodology – ADP, SHY; Formal analysis – ADP; Investigation – ADP, SHY; Writing – Original Draft – ADP; Review and Editing – SHY, PE, ADP, WLL; Supervision – SHY, PE, WLL; Resources – SHY, WLL; Funding acquisition – PE, SHY

## Ethics approval

This research involved collecting honey bees from forest reserves and national parks. The permit applications were approved by the Perak State Forestry Department to collect the samples from Temenggor forest reserve with the approval number: JH/100 JLD.31 (46), and the Sabah Biodiversity Centre, with the access license JKM/MBS.1000-212 JLD.16 (117) and Sabah Parks approval TTS 100-6/2 JLD.34 issued to SHY.

## Consent to participate

Not applicable

## Consent to publish

Not applicable

## Data availability

All the data including the GenBank accession numbers are available within the paper and its Supplementary Information.

