## Supplementary Information for "Integrative taxonomy and distribution of *Apis* species in Malaysia"

**Supplementary figure**

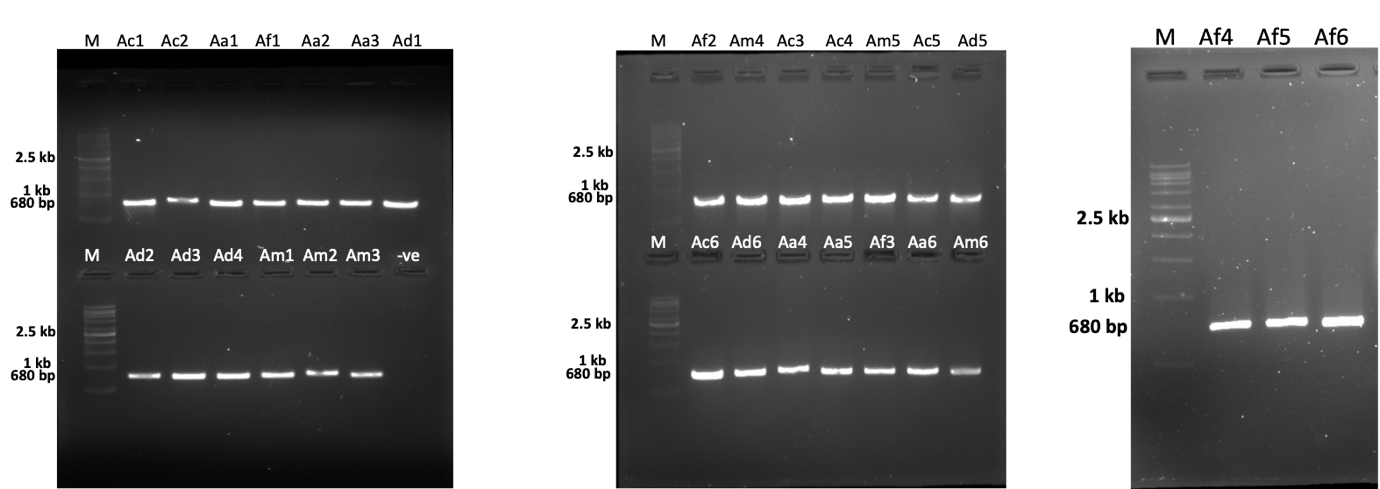

a

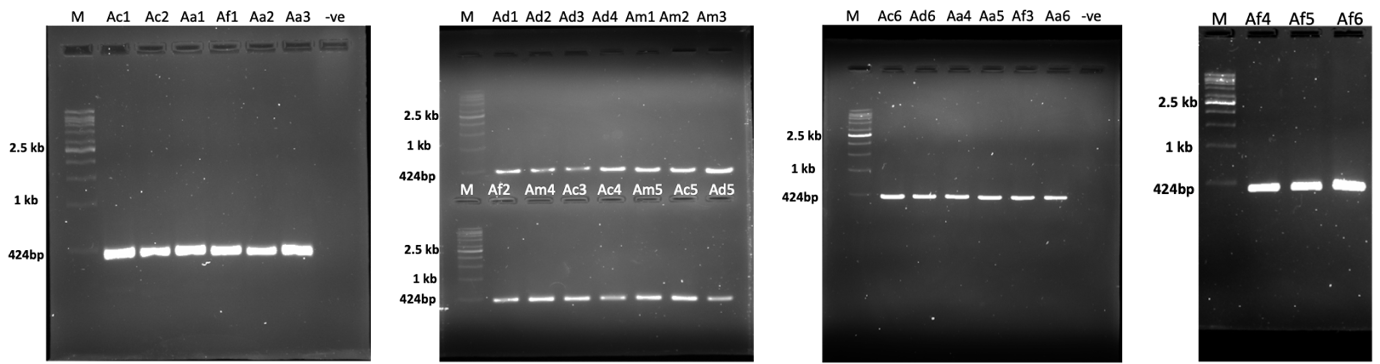

b

**Supplementary Fig S1**: Agarose gel electrophoresis (1.2%) of PCR products from honey bee genomic DNA samples using primers (a) LepF1 & LepR1, and (b) WLO001F-Af & WLO001R-Af with the expected amplicon sizes – 680 bp (CO1 gene segment) and 424 bp (16S rRNA gene segment) respectively. Lanes; 1 kb Ladder: VC 1 kb ladder; lanes denoted by Ac1, Ac2, Ac3, Ac4, Ac5, and Ac6 : *A. cerana* colonies (1-6); Aa1, Aa2, Aa3, Aa4, Aa5, and Aa6: *A. andreniformis* colonies (1-6); Af1, Af2, Af3, Af4, Af5, and Af6: *A. florea* colonies (1-6); Ad1, Ad2, Ad3, Ad4, Ad5, and Ad6: *A. dorsata* colonies (1-6); Am1, Am2, Am3, Am4, Am5, and Am6: *A. mellifera* colonies (1-6); and lanes -ve: no template control.

**Supplementary table**

**Supplementary Table S1** (separate file: Supplementary_TableS1.xlsx)

**Sample collection**

**Permits** (separate file: SampleCollectionPermits.pdf)

**Standard Operating Protocol** (separate file: SampleCollection_SOP.docx)
